# Habitat complexity reduces feeding strength of freshwater predators

**DOI:** 10.1101/2025.02.22.639633

**Authors:** Mireia Aranbarri, Lorea Flores, Ioar de Guzmán, Aitor Larrañaga, Björn C. Rall, Julia Reiss

**Affiliations:** Laboratory of Stream Ecology; Department of Plant Biology and Ecology, Faculty of Science and Technology; University of the Basque Country, UPV/EHU PO Box 644; 48080 Bilbao; Spain; INRAE, UMR 1224, Ecologie Comportementale et Biologie des Populations de Poissons, Aquapôle, quartier Ibarron, 64310 Saint-Pée sur Nivelle, France; Aquatic Ecology and Evolution, Department of Biology, University of Konstanz/Egg, Universitätsstraße 10, 78464 Konstanz, Germany; Centre for Pollution Research and Policy, Brunel University of London, Uxbridge, UB8 3PH, UK

## Abstract

1. The physical structure of an environment potentially influences feeding interactions among organisms, for instance, by providing refuge for prey. We examined how habitat complexity affects the functional feeding response of an ambush predator (damselfly larvae *Ischnura elegans*) and a pursuit predator (backswimmer *Notonecta glauca*) feeding on the isopod *Asellus aquaticus*.

2. We ran experiments in aquatic microcosms with an increasing number of structural elements (0, 2, or 3 rings of plastic plants in different spatial configurations), resulting in five habitat complexity levels. Across these levels, predators were presented with different prey densities to determine the functional response pattern. The experimental design and analysis allowed us to test for effects of structure presence, amount, and complexity level on functional response in one pass, without confounding predictors.

3. The feeding for both predators across all complexity levels was best described by a type II functional response model, and habitat drove feeding strength. Regarding the latter, the predators showed different responses to the complexity treatments. The overall feeding rate of *I. elegans* was mainly explained by the absence vs. presence of structure. Yet, in the case of *N. glauca,* feeding rate was strongly dependent on habitat complexity with the predator showing a unique maximum feeding rate (i.e. the inverse of the handling time) for each complexity level and a decreasing attack rate with increasing amount of habitat.

4. On average, prey consumption by both predators was reduced when complex structures were present, compared to the ‘no habitat structure’ environment (e.g. consumption more than halved for some treatments). Our findings demonstrate that habitat complexity dampens feeding rates and therefore plays a key role in the stability of freshwater ecosystems.

## 1. Introduction

The physical structure of an environment is a crucial factor in shaping community composition and the ecological processes driven by living organisms. Habitat complexity has been shown to influence population density, body size, and species richness of invertebrates in both terrestrial and aquatic ecosystems (Flores et al., 2016; Soukup et al., 2022; White & Walsh, 2020). Furthermore, habitat complexity is also likely to influence the interactions among organisms by providing more refuge, thereby reducing predation risk for prey, especially when prey densities are low (Barrios□O’Neill et al., 2015; Hauzy et al., 2010; Orrock et al., 2013; Vucic-Pestic et al., 2010a,b). Indeed, previous studies have demonstrated that increasing habitat complexity can lower predation pressure on prey, thus reducing the top-down control exerted by predators (Chang & Todd, 2023; Kalinkat et al., 2013; Mocq et al., 2021). Yet habitat complexity could also aid predators in their search for prey, particularly those operating in three-dimensional environments (Pawar et al., 2012), as observed in the case of a freshwater copepod preying on ciliates (Reiss & Schmid-Araya, 2011). Therefore, habitat complexity can potentially modify or shape energy flows through food webs, influencing feeding relationships between resources and consumers and among predators and prey.

Functional response models are a suitable tool when it comes to addressing the effects of habitat complexity on species interactions, as they describe the relationship between prey density and the intake of prey by a predator (Holling, 1959a; Li et al., 2018). More generally, functional response models have been central in understanding interaction strengths, population dynamics and ecosystem stability (Berlow et al., 2009; Kratina et al., 2022; Rall et al., 2008; Williams & Martinez, 2004). Characterizing an organism’s feeding behaviour (in short-term experiments), feeding rate, F, is a function of prey density, N, and determined by the time required for predators to find, attack, capture, handle, and ingest their prey (Holling, 1959a; Li et al., 2018). When observing feeding rate for a range of increasing prey density, then these rates often have a hyperbolic saturating shape, called the type II functional response (Figure 1a). The initial increase in the feeding rate, F, is driven by the attack rate, a, which was originally termed ‘instantaneous rate of discovery’ (Holling, 1959a); and other synonyms include the terms capture-, filtration- or search rate. The handling time, T_*h*_, controls the saturation of the curve (Figure 1a). Alternatively, the function can be written using the maximum feeding rate, *F_*max*_* =- 1/*T_*h*_*, and the half saturation density, *N_*half*_* =- 1/(*aT_*h*_*) - i.e. the prey density where half of the maximum feeding rate is reached (Real, 1977; Figure 1a). Following the ‘Holling approach’ (i.e. models that use attack rate and handling time *sensu* Holling, 1959a) and the ‘Real approach’ (i.e. models using maximum feeding rate and half saturation density *sensu* Real 1977), the full functional response models are

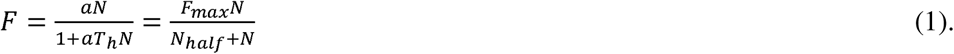

**Figure 1.**
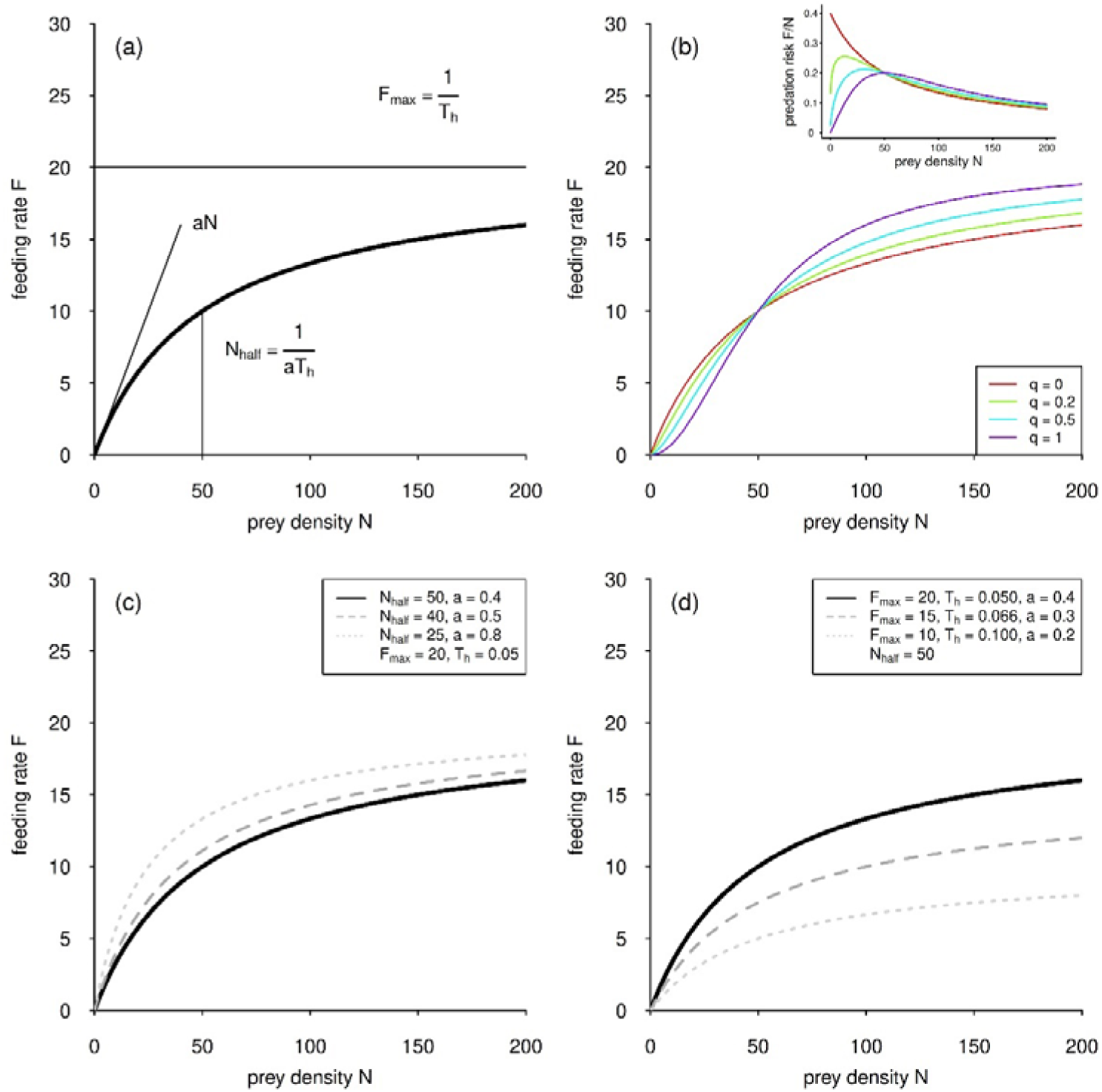
**(a)** The type II functional response (bold line) is either controlled by the parameters attack rate, *a*, and handling time, *T_h_* (Holling 1959b), or the maximum feeding rate *F_max_* and the half saturation density *N_half_* (Real 1977). **(b)** The generalized functional response model allows for a seamless shift between the hyperbolic type II functional response (*q* = 0) and a ‘strict’ type III functional response (*q* = 1). Inlay: The predation risk of a single prey item decreases in the case of a hyperbolic type II functional response (*q* = 0); when the function becomes s-shaped (type III) the predation risk increases with increasing prey density, leading to small but stable populations. **(c)** Example where the Holling and Real models (shown in [a]) are interchangeable. The feeding rate decreases primarily at low prey densities. Although it appears that the maximum feeding rate also declines, this is misleading; in reality, the maximum rate is simply reached at much higher prey densities. **(d)** Example where the models are not interchangeable. The variation in feeding rates as a function of parameter values is clearly visible. The feeding rate decreases uniformly across all prey densities, i.e. the decline is proportional and independent of prey density.

In addition to the hyperbolic shape, other functional response patterns are observed, for instance Holling (1959b) reported an s-shaped functional response for mammal predators under field conditions. The s-shape (type III response) can be observed when both the feeding- and attack rate are a function of prey density, meaning that the predator optimizes hunting prey with increasing prey density. The simplest way to include this feature in the above models is to assume a linear increase in the attack rate, *a =- bN*, with *b* being the attack coefficient (Juliano, 2001). Dubbed the generalized functional response model, this approach allows for a seamless switch from a type II functional response to an s-shaped type III functional response (Figure 1b), promoted by many modelling studies (Jeschke et al., 2004; Kalinkat & Rall, 2015; Rall et al., 2008; Williams & Martinez, 2004). Here, attack rate is a power-law function of prey density (a =- bN^*q*^). If the ‘shape parameter’ q is 0, the functional response is type II, and if the shape parameter q is 1, the functional response is a ‘strict’ type III response (Kalinkat et al., 2023). The generalized functional response models are

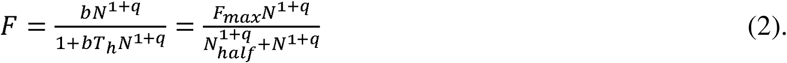

Despite being a commonly reported functional response type (Jeschke et al., 2004; Kalinkat & Rall, 2015) the type II functional response is known to produce unstable population dynamics, especially under environmental enrichment (e.g. Rall et al., 2008; Rosenzweig, 1971). In contrast, the s-shaped functional response is known to stabilize population dynamics and enhance biodiversity, at least in modelling studies (Rall et al., 2008; Williams & Martinez, 2004). The difference of the hyperbolic type II and s-shaped type III functional response can be explained by the differences in predation risk (Figure 1b). The stabilizing effect of the type III functional response can be due to a low predation risk per prey individual (Figure 1b inlay). Generally, prey growth increases more slowly than predation risk, resulting in a low but stable population that remains constrained by top-down pressure (in contrast to a type II response).

The detection of an s-shaped (type III) functional response remains methodologically challenging (Kalinkat et al., 2023) and some researchers argue that apparent type III responses might mainly arise from statistical artefacts rather than from underlying biological mechanisms (DeLong et al., 2025). Undeniably, the experimental design is critical in identifying functional response patterns accurately (DeLong et al., 2025; Sarnelle & Wilson, 2008), and habitat presence might be a particularly influential predictor in functional response experiments (Kalinkat & Rall, 2015). For instance, habitat complexity has frequently been cited as a key driver of type III functional responses (Barrios□O’Neill et al., 2015; Kalinkat et al., 2013; Vucic□Pestic et al., 2010a,b).

Testing how refuge availability and habitat complexity alter trophic relationships requires careful experimental design to avoid confounding structural quantity with habitat complexity (Flores et al., 2016; St. Pierre & Kovalenko, 2014). In a laboratory experiment with a centipede predator and springtail prey, Kalinkat et al. (2013) observed that increasing the amount of structure (adding leaves) provided more surface area for both predator and prey, but did not create refuge. In the latter study, the observed changes in space-dependent functional response parameters did not vary per unit surface, suggesting a dilution effect rather than refuge use (Kalinkat et al., 2013). Importantly, habitat complexity can increase not only by adding more structures, but also through changes in the spatial arrangement of a fixed number of elements (Flores et al., 2016). Furthermore, certain forms of habitat complexity can represent obstacles without contributing to available surface area (Hauzy et al., 2010).

In addition to habitat complexity, the predation strategy of the predator is an important factor determining the interaction strength of predator and prey (Almany, 2004; Schmitz, 2017) and, by extension, the stability of food webs. These strategies include ‘ambush’ (also known as ‘sit-and-wait’ tactics) and ‘pursuit’ (active chasing), and these traits can determine spatial utilization and interactions with environmental structures (Pawar et al., 2012; Schmitz, 2017). For instance, ambush predators, exemplified in aquatic systems by species like dragonfly and damselfly nymphs, occur in complex habitats that provide ample hiding spots, enhancing their ability to remain concealed and ambush prey (Chang & Todd, 2023; Mocq et al., 2021; Soukup et al., 2022). Conversely, pursuit predators, such as predatory fish, certain types of predatory shrimp or backswimmers require more open and less obstructed spaces to effectively chase down their prey. Dense structures within complex habitat, however, can impede their navigation and reduce hunting efficiency (Almany, 2004; Warfe & Barmuta, 2006). Therefore, habitat complexity is likely to have varying impacts on predatory success depending on the predator’s hunting strategy. For instance, in aquatic environments, a more complex habitat may favour ambush predators by providing increased hiding opportunities, while posing challenges for active swimming predators, thereby potentially reducing their hunting efficiency (Almany, 2004; Dunn & Hovel, 2020; Froneman & Cuthbert, 2022). Studying how functional response of different predators is modulated by habitat structure is crucial to understand trophic interactions and energy transfer through real food webs in nature.

We aimed to explore how two different predators with contrasting hunting strategies are affected by habitat complexity in their search and consumption of prey. Specifically, we tested how habitat complexity (measured as [1] the presence of structure, [2] the amount of structure and [3] the spatial arrangement of structure) influenced functional response parameters, addressing three hypotheses as follows.

1. Habitat complexity alters the shape of the functional response curve, as more structure can create refuge for prey and makes detection of prey more challenging, particularly at low prey densities (Figure 1b).
2. Feeding rates generally decrease with increasing amount of habitat structure (Figure 1c), resulting in altered functional response parameters. We expected that attack rate rather than handling time would be affected by habitat complexity meaning feeding rate would be reduced at low prey densities (Figure 1b).
3. The pursuit predator (*N. glauca*) will show a different response to habitat structure compared to the ambush predator (*I. elegans*).

## 2. Material and Methods

### 2.1 Experimental set-up

The experiment was carried out using individual microcosms (9.5 cm diameter and 11.5 cm high circular beakers). The prey (water hog-louse *Asellus aquaticus*) was provided with leaves to feed on. Hence, air-dried alder leaves (*Alnus glutinosa* L.) were weighed, added to each microcosm (2 g each) and conditioned in 500 mL water (1 part of filtered pond water to 5 parts of tap water) for 7 days. The water was renewed when the prey was introduced to the microcosms. It should be noted that the individuals were not able to hide in the leaves, and we therefore ignored the leaves in any estimates of habitat complexity. Every microcosm was served by one air-stone, and it was sealed with cling film to prevent evaporation. The experiment was run in a controlled-temperature room (15 °C) with a 14 h light:10 h dark cycle.

When manipulating habitat complexity, it is important to disentangle the effects of complexity *per se* from the effects of the amount of structure in the microcosm (see Kalinkat et al. 2013 for detailed reasoning). We used plastic rings to create both structure and different levels of complexity (Figure 2). These rings were artificial plastic plants mimicking the aquatic plant *Ceratophyllum* spp. (Code No. FRF 491, Fish are Fun®). Habitat complexity was created in three ways: 1. as the presence vs. absence of habitat structure; 2. as the amount of structure (0, 2 or 3 of plastic plant rings); and 3. as two spatial configurations of structures in the case of 2 and 3 rings (Figure 2; and Flores et al. 2016), resulting in 5 complexity levels. The order of these five levels (from ‘simple’ to ‘complex’ [levels 0, 1, 2, 3 and 4]) was determined by measuring the fractal dimension in the feeding arena (Figure 2; and Flores et al. 2016).

**Figure 2.**
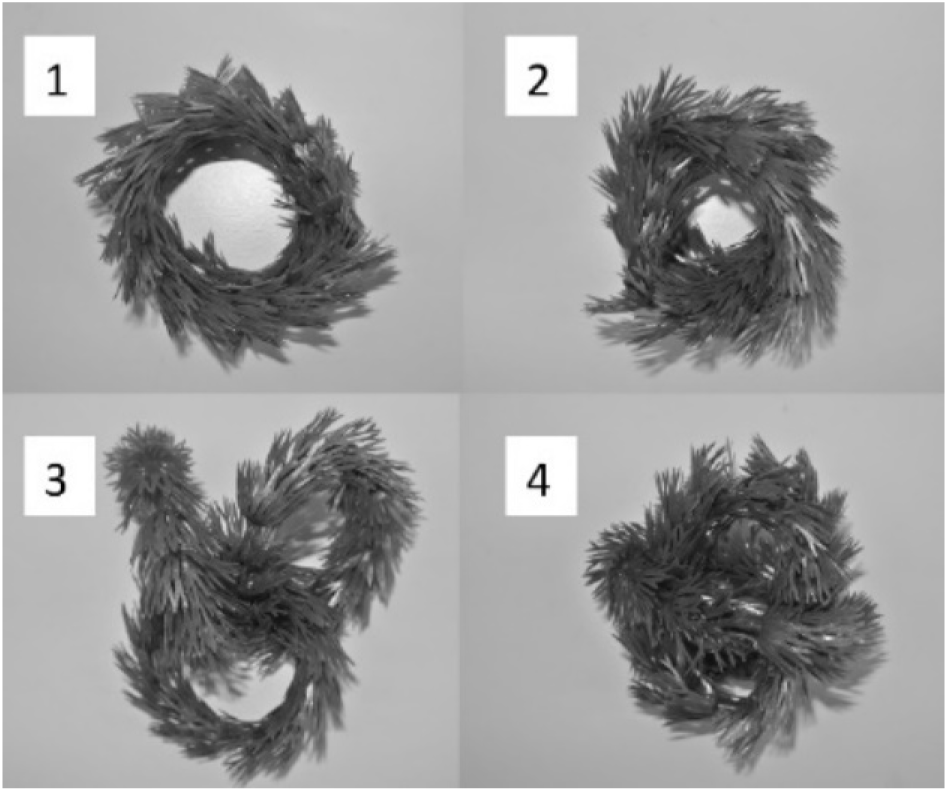
Photographs of the structures used to create habitat complexity in microcosms with ‘structure present’. The basic unit of each structure was a plastic plant strip (mimicking *Ceratophyllum* spp.), joined up as a ring (∼ 8 cm in diameter) and four levels of fractal dimension were created with them: 1) level 1 consisted of two rings aligned, with a fractal dimension (D) of 1.77; 2) level 2 consisted of two rings twisted into each other (D = 1.80); 3) level 3 consisted of three rings locked together (D = 1.81) and 4) level four was a ball made from 3 rings together (D = 1.83). Fractal dimension was used to assign a categorical value for ‘complexity’ to each treatment (zero to four). This design therefore also gave two levels of ‘amount of structure’ - 3g for complexity level 1 and 2 and 4.5 g for complexity level 3 and 4. Figure taken from Flores et al. (2016).

### 2.2 Feeding functional response

We tested the feeding functional response of the ambush predator *Ischnura elegans* (van der Linden) (16.8 mm average length and 3.5 mm average head width) and a pursuit predator *Notonecta glauca* (L.) (14.7 mm average length and 4.9 mm average width) preying on *A. aquaticus* (6.6 mm average body length). Applying body dimension to dry weight regressions from the literature (Baines et al., 2015; Benke et al., 1999; Reiss et al., 2011), we estimated these species to have an average dry weight of 8.6, 10.5 and 2.1 mg per individual respectively.

The predators were able to get into the structures added to the microcosms, which, thus, did not function as a full refuge for prey. The predators were allowed to feed for 24 hours. We used time blocks to run a replicate each, which were all started two days apart from each other because of the time needed to sort *A. aquaticus* individuals. Further, the blocks had the advantage that this approach allowed us to add ‘very high’ density treatments after running the first replicate and to further adjust the number of replicates needed after observing feeding during the first time block. Prey densities for *I. elegans* were 1, 3, 5, 10, 30, 80 and 120 individuals per microcosm. Prey densities for *N. glauca*, were 1, 3, 5, 10, 30, 80, 120 and 180 individuals per microcosm. Replication varied from 1 to 6 replicates per treatment (mostly 3 for *I. elegans* and 6 for *N. glauca*) and in total we ran 297 microcosms. The data is available on GitHub (https://github.com/b-c-r/CRITTERdata) and on Zenodo (Flores et al., 2025).

### 2.3 Data Analysis

Equations 1 and 2 above assume that prey is replenished, however in our laboratory experiments, prey density declined through time. The problem of prey depletion — and its possible solutions — are described by Rosenbaum & Rall (2018). Following the latter authors, we used the Rogers Random Predator Equation (Rogers, 1972; Royama, 1971) modified with the Lambert W function (Bolker, 2008) to fit type II functional responses, simulating prey decline (Rosenbaum & Rall, 2018). For estimating the generalized functional response parameters, we directly fitted simulated time series to our data. These methods prevent biased parameter estimations (see details in Rall et al., 2025a; Rosenbaum & Rall, 2018).

We first investigated if the shape of the functional response was hyperbolic or s-shaped for all ten treatments (5 complexity treatments and 2 predator species). This was achieved by fitting the generalized functional response model (Equation 2, Figure 1b) allowing for a seamless shift between type II and type III functional responses (Rosenbaum & Rall, 2018).

After determining the shape of the functional response, we tested a suite of functional response models to address the effects of habitat on the functional response parameters (see (Rall et al., 2025a). This suite of models tested for 1) the assumption that habitat does not influence feeding, 2) an effect of no structure vs. structure, 3) for an effect caused by the amount of structure and 4) for effects due to the complexity (fractal dimension) of the structure (while being related to ‘amount’). The models targeted two parameters or three parameters (case generalized functional response) at once (Table 1).

**Table 1:**
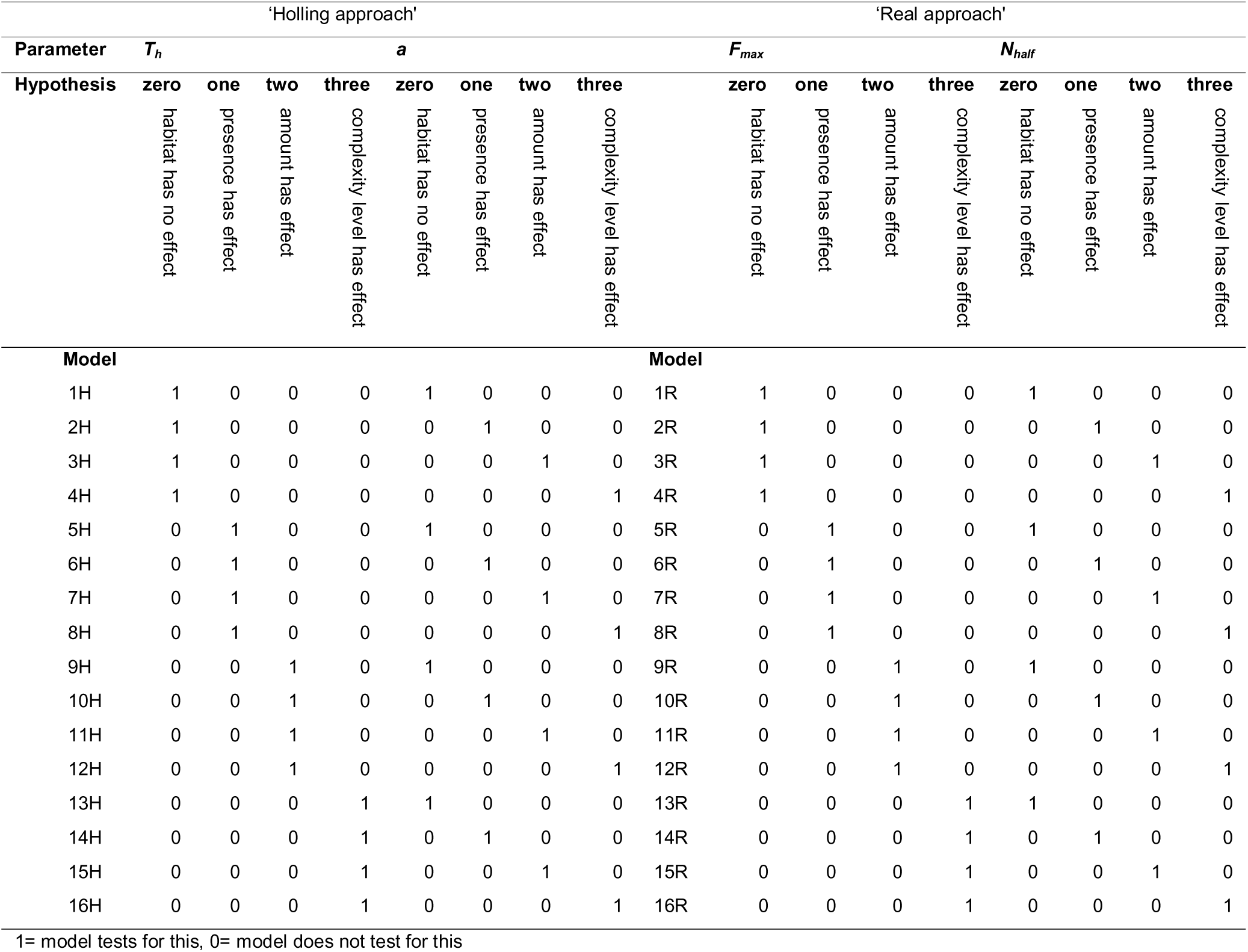
Overview of models tested on two parameters at once to show if feeding rate was affected by habitat complexity. In total, four parameters are tested, yet only two of them in one pass as this reduced the testing from a possible 128 combinations to 32 models. Also, the four parameters correspond to two different simulation models (dubbed ‘H models’ and ‘R models’; after Holling 1959a,b and Real 1977, respectively). For each parameter, there are four hypotheses: ‘Zero’ assumes that habitat has no effect; ‘One’ assumes that presence has an effect (2 levels: present and absent); ‘Two’ assumes that amount has an effect (3 levels: 0, 2 and 3 plastic plant rings) and ‘Three’ assumes that complexity level (fractal dimension) has an effect (5 levels: 0 to 4 complexity levels).

Testing four assumptions on two parameters in a fully factorial fashion gives a suite of 16 models for the ‘Holling’ approach (1H to 16H) and the ‘Real’ approach (1R to 16R) of functional responses each (Table 1). We dubbed the approach using *a* and *T_h_* ‘Holling’ and the approach using *F_max_* and *N_half_* ‘Real’ as these authors were the first to introduce these parameters (Holling, 1959a,b; Real, 1977). Consequently, models 1H to 16H targeted parameters *T_h_*and *a* (*sensu* Holling 1959a,b) and 1R to 16R targeted *F_max_*and *N_half_* (*sensu* Real 1977). In case we detected any type III functional response, the full-factorial scheme would have need 64 models for each functional response formulation (see Rall et al. 2025a).

In brief, the reasoning for fitting both Holling- and Real- approaches to functional response models lies in testing effects of a predictor like habitat complexity. Although Holling- and Real- equations can be generally translated into each other (Figure 1a, Equations 1 and 2), this does not necessarily work when the parameters are connected to an overarching model, like in our case, habitat complexity levels. For instance, as long as both parameters depend on the same habitat complexity measure (e.g. five levels of habitat complexity), the parameter values of the Holling-approach and the Real-approach can be calculated from each other. Yet, in some cases, models are not exchangeable, e.g. if the attack rate / half saturation densities depend on the amount of structural elements and handling time / maximum feeding rate are fitted to the level of complexity. This phenomenon is caused by the fact that the half saturation density is not simply the inverse of attack rate but also depends on the handling time. A change in maximum feeding rate, for instance, automatically leads to a proportional change in attack rate, if the half saturation density is constant (see Figure 1c,d and Rall et al. 2025a for more details).

This approach allowed testing for these different habitat predictors on all functional response parameters in one pass and avoided confounding ‘amount of structure’ with ‘complexity’, based on the rationale developed in Flores et. al. (2016). We compared the models by using both the Akaike’s Information Criterion (AIC) and the Bayesian information criterion (BIC), where the lowest AIC and BIC indicates the best (most parsimonious) model. The output includes a point estimate for the parameter and lower and upper confidence intervals (CIs).

Hence, the functional response models provide information about the overall amount of prey eaten, driven by the maximum feeding rate or its inverse, the handling time. This effect is most obvious at high prey densities, where the curve reaches its asymptote (Figure 1a, c). The attack rate, and the half saturation density predominantly control the feeding rate at lower densities (Figure 1a, d), and we were able to compare those parameters between habitat treatments.

We provide the underlying code on GitHub (https://github.com/b-c-r/CRITTERcode) and as a citable version on Zenodo (Rall, et al., 2025b). This includes a README for the description of the code. The latter also contain detailed theoretical and statistical considerations, and methods in the supplement.

## 3. Results

The shape of the functional response curves for both *N. glauca* and *I. elegans* (Figures 3 and 4) consistently fit best to a type II model for all five complexity levels, as established by fitting the generalized functional response model to the data (Table 2). Applying this model to both species and all complexity levels, the shape parameter *q* was never significantly different from zero, hence feeding rates followed a functional response type II (Table 2).

**Figure 3.**
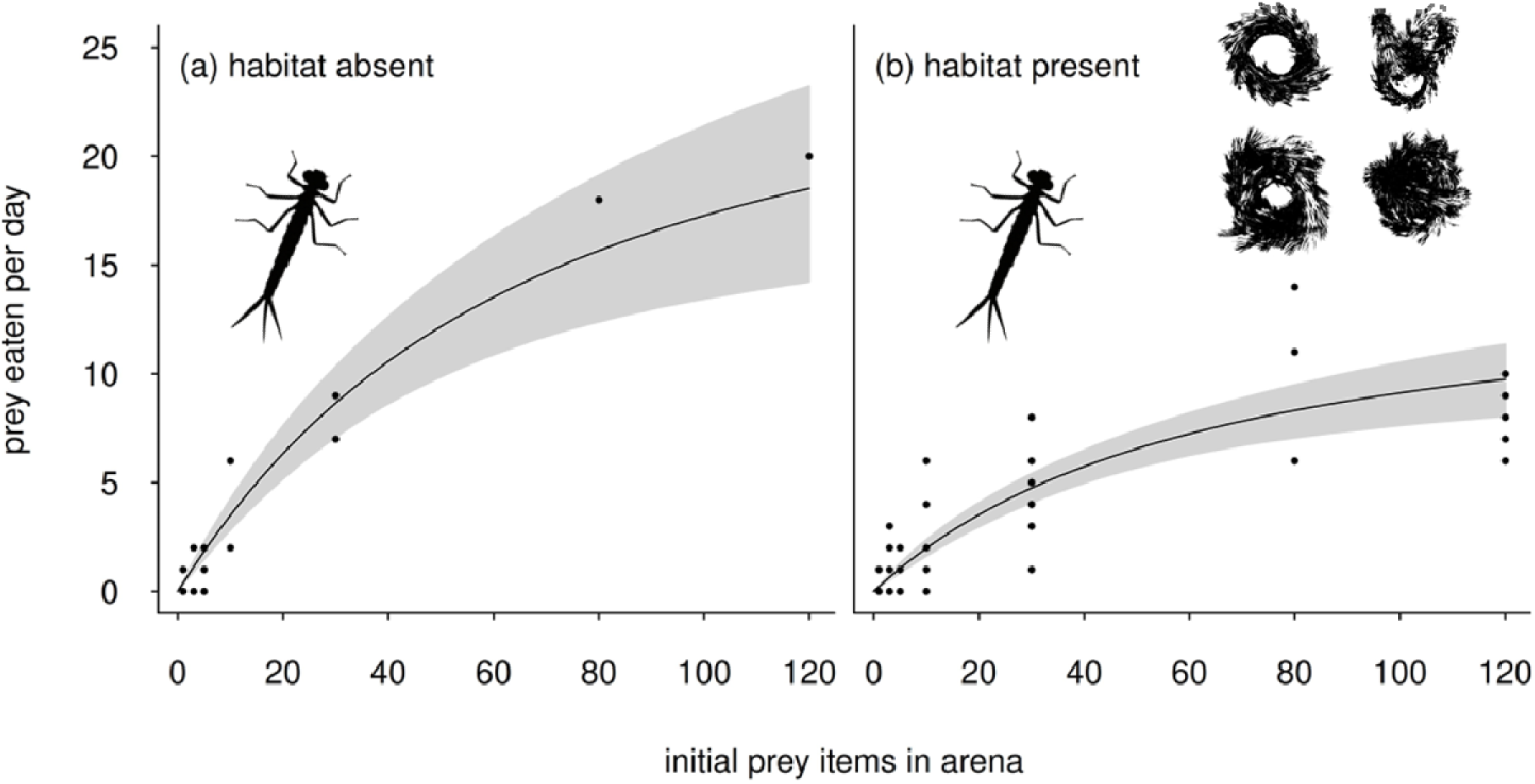
Functional response curves for the ambush predator *I. elegans* feeding in microcosms (arena) without structure **(a)** and in arenas with plastic plant structure **(b)**. The right panel (b) shows data across four habitat complexity levels because the most significant statistical model for the *I. elegans* feeding pattern was the difference between complexity level zero (no structure) and treatments with structure present. Grey shaded areas denote the 95% confidence intervals.

**Figure 4.**
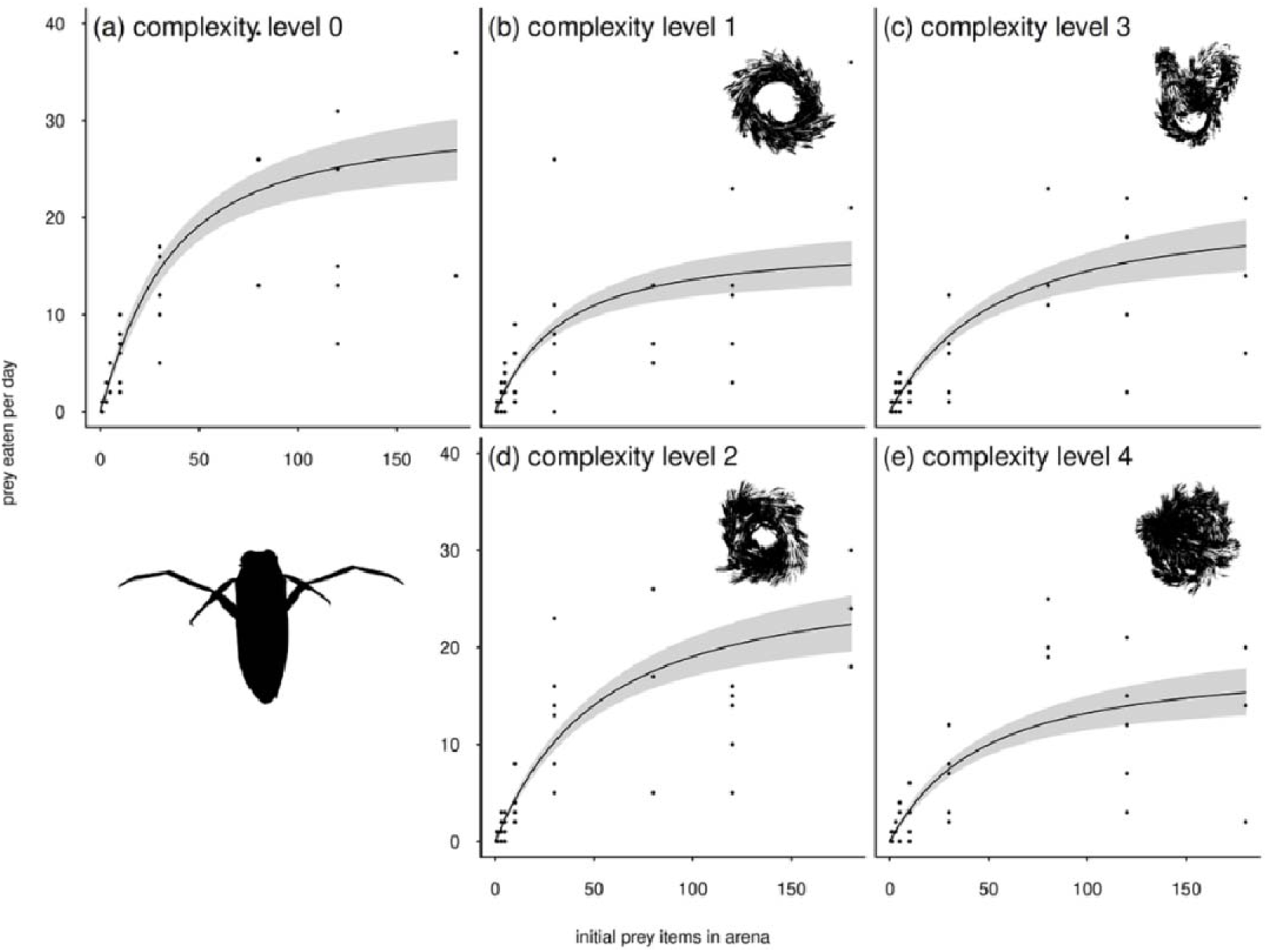
Functional response curves for the pursuit predator *N. glauca* in feeding arenas with different complexity level created with plastic plant rings (see images in panels b to e and Figure 2). Habitat complexity levels 1 and 2 (b, d) were created with two rings of plastic plants, while levels 3 and 4 (c, e) had three rings of plastic plants. All five habitat complexity levels are shown (and not summarised in one panel as in Figure 3) because the most significant statistical model for the *N. glauca* feeding pattern assumed that each complexity level has a unique handling time, and that attack rate decreases with increasing amount of habitat structure. Grey shaded areas denote the 95% confidence intervals.

**Table 2:** Testing for the functional response type for two predators feeding at 5 complexity levels (0 to 4), by using the generalized functional response model (Rosenbaum & Rall, 2018; Vucic-Pestic et al., 2010b; Williams & Martinez, 2004). We observed a declining proportion of prey eaten with increasing complexity level; consistent with a type II functional response (indicated by the value for the shape parameter *q*). The shape parameter *q* is not significantly different from zero in all cases; hence the feeding rate follows a functional response type II.

| Predator | Complexity | $q$ | Significance | Type |
| --- | --- | --- | --- | --- |
| *Ischnura elegans* | 0 | 0.116 | n.s. | II |
| *Ischnura elegans* | 1 | 0.047 | n.s. | II |
| *Ischnura elegans* | 2 | -0.001 | n.s. | II |
| *Ischnura elegans* | 3 | -0.001 | n.s. | II |
| *Ischnura elegans* | 4 | 0.193 | n.s. | II |
| *Notonecta glauca* | 0 | -0.011 | n.s. | II |
| *Notonecta glauca* | 1 | -0.006 | n.s. | II |
| *Notonecta glauca* | 2 | 0.194 | n.s. | II |
| *Notonecta glauca* | 3 | -0.002 | n.s. | II |
| *Notonecta glauca* | 4 | -0.001 | n.s. | II |

Therefore, we tested a total of 32 type II functional response models (Table 1) to test for the effect of habitat. For the functional response exhibited by *I. elegans,* the best fitting model (both lowest AIC and BIC, Table 3) was ‘model 5R’ (Table 3, Figure 3). This model assumed that the effect of structure presences vs. absence overrides all other predictors. In the ‘5R’ model, *F_max_* takes a value of 28 prey eaten in 24 h when no structure is present (CI: 19.4 to 40.8). However, when structure was present, then *F_max_* was 14.6 (CI: 10.7 to 29). *N_half_* was not affected by the habitat and takes the value of 57 prey items per arena (CI: 34 to 94).

**Table 3:** The six best models (see Table 1 for explanation of each model) that describe the functional response parameters *T_h_* and *a* (model names includes an ‘H’), or *F_max_*and *N_half_* (model names includes an ‘R’) and for predators *I. elegance* and *N. glauca* feeding on *A. aquaticus*. The full suite of models can be found in Rall et al. (2025a). The best fit of the models was tested using AIC (AIC generally performs better for models that have more parameters) and BIC. A low value indicates the best fit.

| AIC<br>Model | df | dAIC | BIC<br>Model | df | dBIC |
| --- | --- | --- | --- | --- | --- |
| ***Ischnura elegans*** |  |  |  |  |  |
| **Model 5R** | **3** | **0.000** | **Model 5R** | **3** | **0.000** |
| Model 7R | 4 | 0.265 | Model 5H | 3 | 1.583 |
| Model 6H | 4 | 0.952 | Model 7R | 4 | 2.731 |
| Model 6R | 4 | 0.952 | Model 6H | 4 | 3.418 |
| Model 5H | 3 | 1.583 | Model 6R | 4 | 3.418 |
| Model 13R | 6 | 2.107 | Model 7H | 4 | 4.718 |
| ***Notonecta glauca*** |  |  |  |  |  |
| **Model 15H** | **7** | **0.000** | **Model 15H** | **7** | **0.000** |
| Model 16H | 10 | 1.274 | Model 9R | 3 | 1.004 |
| Model 16R | 10 | 1.274 | Model 11H | 4 | 1.485 |
| Model 15R | 7 | 1.990 | Model 11R | 4 | 1.485 |
| Model 14R | 7 | 4.189 | Model 13R | 6 | 1.507 |
| Model 13R | 6 | 4.854 | Model 15R | 7 | 1.990 |
| Model 12H | 7 | 8.352 | Model 7H | 4 | 2.516 |

The best fitting (most parsimonious) model (both lowest AIC and BIC) for *N. glauca* was ‘model 15H’ (Table 3, Figure 4) and this model assumed that each complexity level has a unique handling time, *T_h_*, and that attack rate, *a*, decreases with increasing amount of habitat structure. Hence, the parameters that influence feeding rate are more complex in this case. Maximum feeding rate (estimated from the handling time [F_*max*_ =- 1/*T_h_*]) was 31 (CI: 26 to 35) prey eaten in 24 h for complexity level 0 (no structure), 18 (CI: 15 to 21) for level 1, 28 (CI: 24 to 33) for level 2, 21 (CI: 17 to 27) for level 3 and 19 (CI: 15 to 23) for level 4 (Figure 4).

The attack rate of *N. glauca* (Figure 4) decreased with increasing number of habitat structural elements (rings, *log*_*10*_(*a*) =- *a*_*intercept*_ + *a*_*slope*_*N*_*rings*_), indicated by a negative slope (-0.144, CI: -0.191 to -0.097).

The reason this more complex model was significant for *N. glauca* over the very similar, but simpler, ‘presence of structure’ model, was that ‘unexpected’ responses were observed when two plastic plant rings had been added to the microcosms, both in terms of feeding- and attack rate (Figure 5). A possible explanation is that the configuration of two rings into two different shapes creates very different habitat types and that this is not the case when three rings were used (the latter essentially created an ideal hiding spot in any case, a ‘dense ball’). For two rings, it is likely that one of the configurations (complexity level 2, two twisted rings, see Figure 2, panel 2) left more space for the agile pursuit predator *N. glauca* to navigate in (Figures 2 and 5).

**Figure 5.**
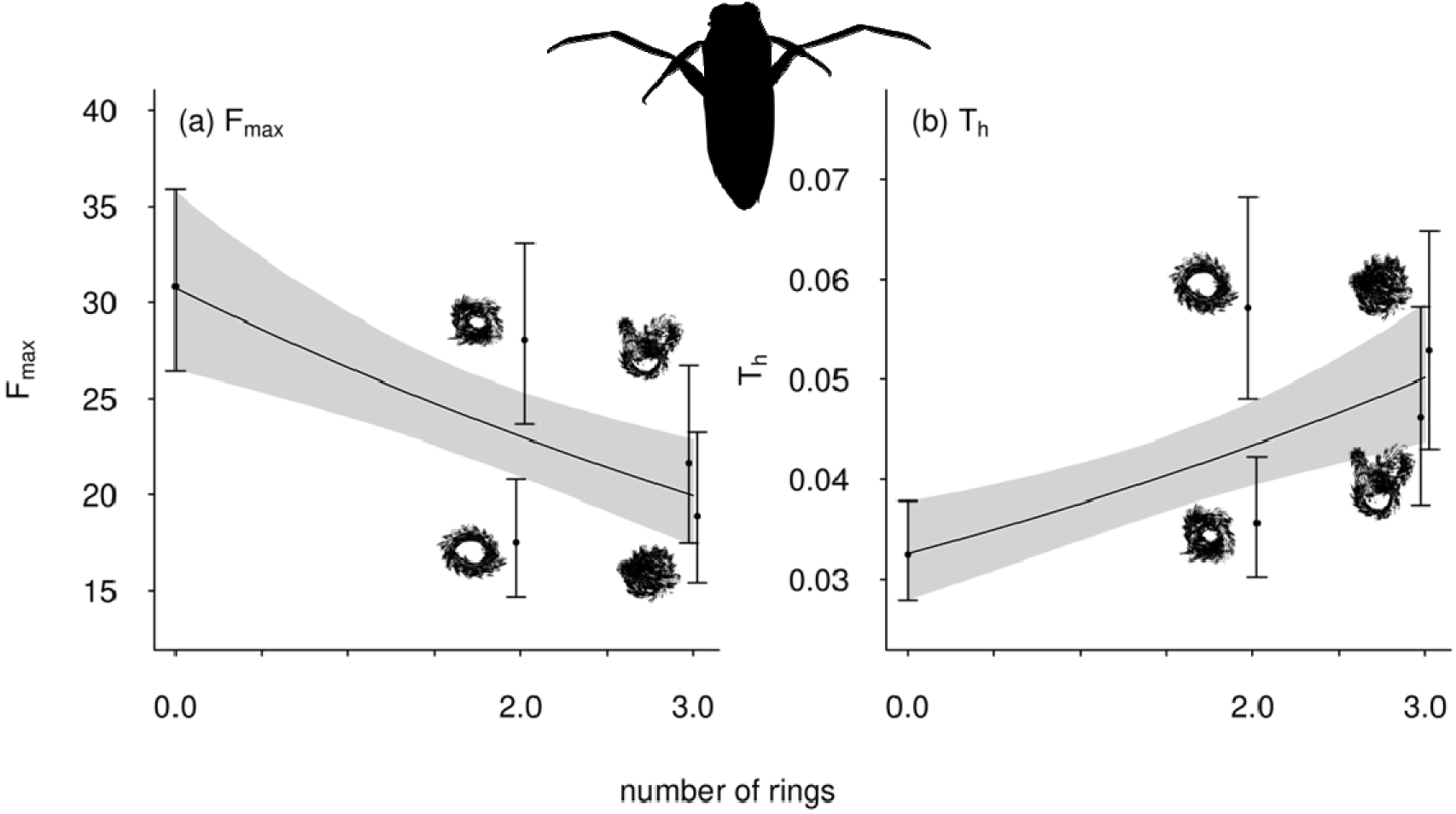
The functional feeding response parameters (**a)** *F_max_* and (**b)** *T_h_* of *N. glauca* feeding on *A. aquaticus*. The amount of structure (number of plastic plant rings) is regressed against maximum feeding rate (a) and handling time (b). When two rings were added, there was an ‘unexpected response’ (high parameter values for complexity level 2) - explaining why a more complex model that assumed an effect of complexity level (levels zero to 4) is more parsimonious compared to assuming that ‘amount of structure’ solely drives the response. Grey shaded areas denote the 95% confidence intervals.

Overall, fitting these models showed that prey consumption was considerably less for both predators when complex structures were present, compared to the no habitat structure environment (Figures 3 and 4). For instance, the maximum number of prey consumed was less than half in some of the ‘no structure’ vs. the ‘structure’ treatments (Figures 3 and 4). To illustrate this difference, in ‘no structure microcosms’, and at a prey density of 120 individuals, the maximum number of prey consumed was 20 and 49 for *I. elegans* and *N. glauca* respectively. In contrast, at the same prey density, *I. elegans* and *N. glauca* respectively fed on a maximum of 9 and 20 *A. aquaticus* in at least one of the structure treatments (the latter were different treatments for the two species, see Figures 3 and 4).

## 4. Discussion

Our findings suggest that habitat complexity affects predator-prey interactions in freshwater ecosystems. Results from our feeding experiments showed that feeding, i.e. the functional response type, did not significantly vary with habitat complexity (all feeding was consistent with a type II response). However, we found support for the hypothesis that increased habitat complexity would decrease feeding rates overall. Both the ambush predator (damselfly larvae *Ischnura elegans*) and the pursuit predator (backswimmer *Notonecta glauca*) showed higher feeding rates in microcosms without structures compared to those with structure. Weighing 8.6 and 10.5 mg (dry weight, dw, per individual) respectively, *I. elegans* and *N. glauca* fed on *A. aquaticus* individuals weighing 2.1 mg dw each. For example, at a prey density of 120 prey, average consumption reached over four (*I. elegans*) and eight (*N. glauca*) times of predator body weight in 24 hours (20 and 33 prey eaten on average by *I. elegans* and *N. glauca* respectively). The latter assumes that prey was ingested whole (which was not the case for *N. glauca*, which left prey body parts after feeding). Taken together, this suggests that complexity affected energy flow considerably and that, while habitat complexity reduced feeding rates by affecting *F_max_*, attack rate and handling time, the overall mechanisms (shape of the functional response curve) remained consistent across different complexity levels.

Generally, it is known that complex structures offer refuge for prey, thereby reducing the efficiency of predators (Froneman & Cuthbert, 2022; Hauzy et al., 2010; Mocq et al., 2021). This reduction in feeding rates due to increased habitat complexity implies a decrease in the top-down control of predators on prey populations (Chang & Todd, 2023). In our study, the reduction in feeding rates was associated with both a reduction in attack rate and an increase in handling time (i.e. an increase in maximum feeding rate while the half saturation density is relatively constant). However, the magnitude of these changes varied across different complexity levels. Previous studies, conducted using microcosm approaches, have shown that attack rate and handling time can respond differently to habitat complexity. For instance, a study on the zooplankton prey *Aracartia longipatella* and the predator *Mesopodopsis wooldridgei* reports strong complexity effects on attack rates (Froneman & Cuthbert, 2022). Yet, the dragonfly larvae *Aeshna cyanea* preying on *Chaoborus obscuripes* larvae was shown to change handling time with complexity (Mocq et al., 2021). Soil mites preying on collembolans displayed varying response in both attack rate and handling time with habitat structure (Hauzy et al., 2010).

Overall, these previous studies also demonstrate that whether habitat is perceived as ‘complex’ by the organisms is context-dependent and that the size proportions of predator to prey – and to habitat – are a key factor. For instance, in feeding experiments with the predatory nematode *Prionchulus muscorum* addition of structure had little effect as predator and prey had the same body shape and mobility - hence the feeding pattern was mainly driven by prey size (Kreuzinger-Janik et al., 2019). Estimating fractal dimension to assign an ‘objective’ complexity level (i.e. a score from low to high) proved valuable for our experiment (see also Flores et al. 2016) but it was obvious that one of the predators (*N. glauca*) was more successful than expected in one case because the structure had much open space, despite its high fractal dimension. Therefore, finding a metric for habitat complexity should include spatial arrangement and consider the dimensions of prey, predator and habitat structure. Clearly, it is important to add this form of realism in laboratory experiments as abiotic factors, such as habitat structure (e.g. Flores et al. 2016) or temperature (e.g. Sentis et al., 2022), can change the outcomes of feeding experiments. More complex physical structure could have a positive effect on overall performance of an assemblage (Flores et al., 2016) because species can feed in their optimum environment (in analogy to their optimum temperature) while predation is dampened. The amount or arrangement of physical structure could, therefore, influence consumer effects and could be an important predictor to consider in ecological research that focuses on energy transfer and ecosystem functioning (Flores et al., 2016).

We hypothesized that at lowest habitat complexity (i.e. microcosms without added plastic plant structures), the shape of the functional response would best fit to a type II functional response because feeding rates would be mainly limited by the time required by the predator to kill and eat its prey. With increasing habitat complexity levels, we expected a sigmoid functional response (i.e. type III functional response), as at low prey densities the efficiency of the predator would be limited by the structures offering multiple hiding places (Kalinkat et al., 2013; Vucic-Pestic et al., 2010a,b). (However, as prey density increases and the amount of refuge becomes saturated, the efficiency of the predator would again increase at intermediate prey densities (Figure 1b,c).

Yet, functional response of both predators was consistently best described by a type II model across all the levels of habitat complexity. Mikolajewski et al. (2015) highlighted a phenotypic plasticity for *Ishnura*, allowing it to adapt to environmental changes while maintaining a type II functional response. Moreover, our design of the habitat structure allowed both predators to enter the potential prey refuge, which may also be a reason for not finding a type III response. One prey per predator was the lowest density in our experiment, which corresponds to a density of 140 individuals per square metre, but in nature, lower abundances, can be observed for many taxa (Larrañaga et al., 2009). Indeed, not considering low prey in laboratory set-ups can hinder the detection of a type III response (Sarnelle & Wilson, 2008). Although the prevalence of type III functional response has been postulated as the way to maintain stability of prey populations and high diversity (Kalinkat et al., 2023; Murdoch & Oaten, 1975; Rall et al., 2008), type II is still the most common functional response in feeding studies (DeLong et al., 2025; Dunn & Hovel, 2020; Kalinkat & Rall, 2015). Even though a type III functional response was not supported by our data, we saw a hint that the interaction of both predators and its prey was stabilizing as feeding rates generally decreased with the presence and with increasing complexity of the habitat structure. The general decrease in interaction strength is associated with more stable population dynamics and higher species richness in model food webs (May, 1972; Rall et al., 2008; Yodzis & Innes, 1992).

The two predators in our experiment differed in terms of their predatory strategies, and the functional response parameters highlighted some of those differences. The pursuit predator (the backswimmer *N. glauca*) exhibited three times the attack rate of the ambush predator (the damselfly *I. elegans*), although a similar handling time, when no structures were present in their environment. Furthermore, as an open-water hunter, *N. glauca* is more likely to operate in a 3D environment (Pawar et al., 2012) compared to *I. elegans* that sits within or on the habitat surface. Indeed, dimensionality of consumer search space is probably a major driver of species coexistence, and the stability and abundance of populations (Pawar et al. 2012).

As predicted, the effect of habitat complexity was stronger for the pursuit predator than the ambush predator. An explanation is the barrier effect that the structures can create for the pursuit predator, reducing the visibility of prey and the capture success (Grabowski & Kimbro, 2005). In agreement with Svensson et al. (2000), habitat heterogeneity significantly reduced predation efficiency in *N. glauca*, highlighting the vulnerability of pursuit predators to habitat complexity. Previous studies have also shown that *Notonecta* spp. are more successful in open environments (Gittelman, 1974), and that factors such as water depth and refuge availability influence its hunting behaviour (Cockrell, 1984).

Our short-term experiment with three species certainly demonstrated that habitat complexity is a key factor to be considered when it comes to trophic interactions. Both predators were more successful in environments without structure and were able to feed on substantially more prey in this open environment. We conclude that habitat complexity has the potential to shape species interactions and carbon flow within the food web. The results of this study demonstrate that increased habitat complexity significantly reduces feeding rates in both *I*. *elegans* and *N. glauca*. The reduction in predation efficiency, driven by decreases in attack rates and increases in handling times, supports the idea that structural complexity provides refuge for prey, limiting predator success. Despite these changes in feeding rates, both species maintained a type II functional response across all levels of habitat complexity, indicating that while habitat structure affects predator efficiency, it does not fundamentally alter the shape of the functional response curve. Overall, these findings underscore the critical role of habitat complexity in shaping trophic interactions and sustaining energy flow. In more complex, natural ecosystems, habitat complexity will drive species coexistence, stability of food webs and diversity of communities.

## Acknowledgements

MA was funded by the Investigo Program funded by the NextGenerationEU initiative, LF was funded by a grant by the Spanish Ministry of Education and Culture, IdG was funded by the Spanish Ministry of Science, Innovation and Universities (TED2021-129966B-C31). JR was supported by a Royal Society of London Starting Grant. Björn C. Rall gratefully acknowledges the funding by the German Science Foundation (DFG) to the Research Unit DynaSym (FOR 5064). We thank the WWT London for the permission to sample damselflies and backswimmers on their grounds and Szymon Szary for his assistance in the field and laboratory. We further thank Arturo Elosegi for his feedback on earlier drafts of this manuscript.

## Author Contributions

Julia Reiss and Lorea Flores conceived the ideas and designed the methodology; Lorea Flores collected the data; Björn C. Rall, Mireia Aranbarri and Ioar de Guzmán analysed the data; Mireia Aranbarri, Björn C. Rall and Julia Reiss led the writing of the manuscript. All authors contributed critically to the drafts and gave final approval for publication.

## Statement on inclusion

This experiment was conducted in the UK, by a German and Spanish research team.

## Data availability statement

Our data, code and statistical methods are available on GitHub (https://github.com/b-c-r/CRITTERdata, https://github.com/b-c-r/CRITTERcode, and https://github.com/b-c-r/CRITTERstatistics) and as citable versions on Zenodo (https://doi.org/10.5281/zenodo.15348769, https://doi.org/10.5281/zenodo.15346225, https://doi.org/10.5281/zenodo.15348995).

## References

Almany, G. R. (2004). Does increased habitat complexity reduce predation and competition in coral reef fish assemblages? Oikos, 106(2), 275–284. 10.1111/j.0030-1299.2004.13193.x

Baines, C. B., McCauley, S. J., & Rowe, L. (2015). Dispersal depends on body condition and predation risk in the semilaquatic insect, *Notonecta undulata*. Ecology and Evolution, 5(12), 2307–2316. 10.1002/ece3.1508

Barriosl□’Neill, D., Dick, J. T. A., Emmerson, M. C., Ricciardi, A., & MacIsaac, H. J. (2015). Predator□free space, functional responses and biological invasions. Functional Ecology, 29(3), 377–384. 10.1111/1365-2435.12347

Benke, A. C., Huryn, A. D., Smock, L. A., & Wallace, J. B. (1999). Length-mass relationships for freshwater macroinvertebrates in North America with particular reference to the southeastern United States. Journal of the North American Benthological Society, 18(3), 308–343. 10.2307/1468447

Berlow, E. L., Dunne, J. A., Martinez, N. D., Stark, P. B., Williams, R. J., & Brose, U. (2009). Simple prediction of interaction strengths in complex food webs. Proceedings of the National Academy of Sciences, 106(1), 187–191. 10.1073/pnas.0806823106

Bolker, B. M. (2008). Ecological models and data in R. In Ecological models and data in R. Princeton University Press.

Chang, C., & Todd, P. A. (2023). Reduced predation pressure as a potential driver of prey diversity and abundance in complex habitats. npj Biodiversity, 2(1). 10.1038/s44185-022-00007-x

Cockrell, B. J. (1984). Effects of water depth on choice of spatially separated prey by *Notonecta glauca* L. Oecologia, 62(2), 256–261. 10.1007/BF00379023

DeLong, J. P., Coblentz, K. E., & Uiterwaal, S. F. (2025). Are type 3 functional responses just statistical apparitions? Ecosphere, 16(4), e70247. 10.1002/ecs2.70247

Dunn, R. P., & Hovel, K. A. (2020). Predator type influences the frequency of functional responses to prey in marine habitats. Biology Letters, 16(1), 20190758. 10.1098/rsbl.2019.0758

Flores, L., Bailey, R. A., Elosegi, A., Larrañaga, A., & Reiss, J. (2016). Habitat complexity in aquatic microcosms affects processes driven by detritivores. PLoS One, 11(11), e0165065. 10.1371/journal.pone.0165065

Flores, L., Reiss, J., Larrañaga, A., Rall, B. C., Aranbarri, M., & de Guzmán, I. (2025). Complexity reduces feeding strength of freshwater predators (CRITTER) - Data(v0.1.2) (Version 0.1.2) [Dataset]. Zenodo. 10.5281/zenodo.15348769

Froneman, P. W., & Cuthbert, R. N. (2022). Habitat complexity alters predator-prey interactions in a shallow water ecosystem. Diversity, 14(6). 10.3390/d14060431

Gittelman, S. H. (1974). Locomotion and predatory strategy in backswimmers (Hemiptera: Notonectidae). American Midland Naturalist, 496–500. 10.2307/2424316

Grabowski, J. H., & Kimbro, D. L. (2005). Predator-avoidance behavior extends trophic cascades to refuge habitats. Ecology, 86(5), 1312–1319. 10.1890/04-1216

Hauzy, C., Tully, T., Spataro, T., Paul, G., & Arditi, R. (2010). Spatial heterogeneity and functional response: an experiment in microcosms with varying obstacle densities. Oecologia, 163, 625–636. 10.1007/s00442-010-1585-5

Holling, C. S. (1959a). Some characteristics of simple types of predation and parasitism. The Canadian Entomologist, 91(7), 385–398. 10.4039/Ent91385-7

Holling, C. S. (1959b). The components of predation as revealed by a study of small-mammal predation of the European Pine Sawfly. The Canadian Entomologist, 91(5), 293–320. 10.4039/Ent91293-5

Jeschke, J. M., Kopp, M., & Tollrian, R. (2004). Consumer-food systems: why type I functional responses are exclusive to filter feeders. Biological Reviews, 79(2), 337–349. 10.1017/S1464793103006286

Juliano, S. A. (2001). Nonlinear curve fitting: predation and functional response curves. In S. M. Scheiner & J. Gurevitch (Eds.), Design and analysis of ecological experiments (2nd ed., pp. 178–196). Oxford University Press.

Kalinkat, G., Brose, U., & Rall, B. C. (2013). Habitat structure alters top-down control in litter communities. Oecologia, 172(3), 877–887. 10.1007/s00442-012-2530-6

Kalinkat, G., & Rall, B. C. (2015). Effects of climate change on the interactions between insect pests and their natural enemies. Climate Change and Insect Pests, 74–91. 10.1079/9781780643786.0074

Kalinkat, G., Rall, B. C., Uiterwaal, S. F., & Uszko, W. (2023). Empirical evidence of type III functional responses and why it remains rare. Frontiers in Ecology and Evolution, 11, 1033818. 10.3389/fevo.2023.1033818

Kratina, P., Rosenbaum, B., Gallo, B., Horas, E. L., & O’Gorman, E. J. (2022). The combined effects of warming and body size on the stability of predator-prey interactions. Frontiers in Ecology and Evolution, 9, 772078. 10.3389/fevo.2021.772078

Kreuzinger-Janik, B., Brüchner- Hüttemann, H., & Traunspurger, W. (2019). Effect of prey size and structural complexity on the functional response in a nematode- nematode system. Scientific Reports 2019 *9*:1, *9*(1), 1–8. 10.1038/s41598-019-42213-x

Larrañaga, A., Basaguren, A., Elosegi, A., & Pozo, J. (2009). Impacts of *Eucalyptus globulus* plantations on Atlantic streams: change s in invertebrate density and shredder traits. Fundamental and Applied Limnology, 175(2), 151–160. 10.1127/1863-9135/2009/0175-0151

Li, Y., Rall, B. C., & Kalinkat, G. (2018). Experimental duration and predator satiation levels systematically affect functional response parameters. Oikos, 127(4), 590–598. 10.1111/OIK.04479

May, R. M. (1972). Will a large complex system be stable? Nature, 238(5364), 413–414. 10.1038/238413a0

Mikolajewski, D. J., Conrad, A., & Joop, G. (2015). Behaviour and body size: plasticity and genotypic diversity in larval *Ischnura elegans* as a response to predators (Odonata: Coenagrionidae). International Journal of Odonatology, 18(1), 31–44. 10.1080/13887890.2015.1012653

Mocq, J., Soukup, P. R., Näslund, J., & Boukal, D. S. (2021). Disentangling the nonlinear effects of habitat complexity on functional responses. Journal of Animal Ecology, 90(6), 1525–1537. 10.1111/1365-2656.13473

Murdoch, W. W., & Oaten, A. (1975). Predation and population stability. In Advances in ecological research (Vol. 9, pp. 1–131). Elsevier. 10.1016/S0065-2504(08)60288-3

Orrock, J. L., Preisser, E. L., Grabowski, J. H., & Trussell, G. C. (2013). The cost of safety: refuges increase the impact of predation risk in aquatic systems. Ecology, 94(3), 573–579. 10.1890/12-0502.1

Pawar, S., Dell, A. I., & Savage, V. M. (2012). Dimensionality of consumer search space drives trophic interaction strengths. Nature, 486(7404), 485–489. 10.1038/nature11131

Rall, B. C., Aranbarri, M., Flores, L., de Guzmán, I., Larrañaga, A., & Reiss, J. (2025a). Complexity reduces feeding strength of freshwater predators (CRITTER) - Supplemental Statistics Report (v.0.1.2). Zenodo. 10.5281/zenodo.15348995

Rall, B. C., Aranbarri, M., Lorea Flores, de Guzman, I., Aitor Larrañaga, & Reiss, J. (2025b). Habitat complexity reduces feeding strength of freshwater predators (CRITTER) —Code (v0.1.2) (Version 0.1.2) [R]. Zenodo. 10.5281/zenodo.15346225

Rall, B. C., Guill, C., & Brose, U. (2008). Foodlweb connectance and predator interference dampen the paradox of enrichment. Oikos, 117(2), 202–213. 10.1111/j.2007.0030-1299.15491.x

Real, L. A. (1977). The kinetics of functional response. The American Naturalist, 111(978), 289–300. 10.1086/283161

Reiss, J., Bailey, R. A., Perkins, D. M., Pluchinotta, A., & Woodward, G. (2011). Testing effects of consumer richness, evenness and body size on ecosystem functioning. Journal of Animal Ecology, 80(6), 1145–1154. 10.1111/j.1365-2656.2011.01857.x

Reiss, J., & Schmid-Araya, J. M. (2011). Feeding response of a benthic copepod to ciliate prey type, prey concentration and habitat complexity. Freshwater Biology, 56(8), 1519– 1530. 10.1111/j.1365-2427.2011.02590.x

Rogers, D. (1972). Random search and insect population models. The Journal of Animal Ecology, 369–383. 10.2307/3474

Rosenbaum, B., & Rall, B. C. (2018). Fitting functional responses: Direct parameter estimation by simulating differential equations. Methods in Ecology and Evolution, 9(10), 2076–2090. 10.1111/2041-210X.13039

Rosenzweig, M. L. (1971). Paradox of enrichment: destabilization of exploitation ecosystems in ecological time. Science, 171(3969), 385–387. 10.1126/science.171.3969.385

Royama, T. (1971). A comparative study of models for predation and parasitism. Population Ecology, 13, 1–91. 10.1007/BF02511547

Sarnelle, O., & Wilson, A. E. (2008). Type III functional response in *Daphnia*. Ecology, 89(6), 1723–1732. 10.1890/07-0935.1

Schmitz, O. (2017). Predator and prey functional traits: understanding the adaptive machinery driving predator–prey interactions. F1000Research, 6, 1767. 10.12688/f1000research.11813.1

Sentis, A., Veselý, L., Let, M., Musil, M., Malinovska, V. and Kouba, A., (2022). Shortlterm thermal acclimation modulates predator functional response. Ecology and Evolution, 12(2), p.e8631. 10.1002/ece3.8631

Soukup, P. R., Näslund, J., Höjesjö, J., & Boukal, D. S. (2022). From individuals to communities: Habitat complexity affects all levels of organization in aquatic environments. John Wiley and Sons Inc. 10.1002/wat2.1575

St. Pierre, J. I., & Kovalenko, K. E. (2014). Effect of habitat complexity attributes on species richness. Ecosphere, 5(2), 1–10. 10.1890/ES13-00323.1

Svensson, B. G., Tallmark, B., & Petersson, E. (2000). Habitat heterogeneity, coexistence and habitat utilization in five backswimmer species (Notonecta spp.; Hemiptera, Notonectidae). Aquatic Insects, 22(2), 81–98. 10.1076/0165-0424(200004)22:2;1-P;FT081

Vucic-Pestic, O., Birkhofer, K., Rall, B. C., Scheu, S., & Brose, U. (2010a). Habitat structure and prey aggregation determine the functional response in a soil predator–prey interaction. Pedobiologia, 53(5), 307–312. 10.1016/j.pedobi.2010.02.003

VuciclPestic, O., Rall, B. C., Kalinkat, G., & Brose, U. (2010b). Allometric functional response model: body masses constrain interaction strengths. Journal of Animal Ecology, 79(1), 249–256. 10.1111/j.1365-2656.2009.01622.x

Warfe, D. M., & Barmuta, L. A. (2006). Habitat structural complexity mediates food web dynamics in a freshwater macrophyte community. Oecologia, 150(1), 141–154. 10.1007/s00442-006-0505-1

White, J. Y., & Walsh, C. J. (2020). Catchmentlscale urbanization diminishes effects of habitat complexity on instream macroinvertebrate assemblages. Ecological Applications, 30(8), e02199. 10.1002/eap.2199

Williams, R. J., & Martinez, N. D. (2004). Stabilization of chaotic and non-permanent food-web dynamics. The European Physical Journal B, 38, 297–303. 10.1140/epjb/e2004-00122-1

Yodzis, P., & Innes, S. (1992). Body size and consumer-resource dynamics. The American Naturalist, 139(6), 1151–1175. 10.1086/285380

